# Rational design of mechanically active RNAs: de novo engineering of functional exoribonuclease-resistant RNAs

**DOI:** 10.64898/2026.01.08.698366

**Authors:** Jule Walter, Leonhard Sidl, Katrin Gutenbrunner, Denis Skibinski, Tim Kolberg, Ivo L. Hofacker, Hua-Ting Yao, Mario Mörl, Michael T. Wolfinger

## Abstract

Mechanically active RNAs represent an emerging class of biomolecules whose function derives from resisting molecular forces. Among them, exoribonuclease-resistant RNAs (xr-RNAs) achieve this by folding into a ring-like topology that physically blocks 5’ → 3’ degradation. However, despite years of structural insight, the rational design of such mechanically functional RNA devices has remained elusive. Here, we describe a mechanics-aware RNA design approach that enables *de novo* engineering of functional xrRNAs. We first identify structural determinants of force resistance by perturbing pseudoknot architecture in a model xrRNA and quantifying resulting efficiencies in the stalling of exoribonuclease XRN1. We then implement these rules in a design framework that integrates explicit topological constraints with molecular dynamics-guided optimization. The resulting synthetic xrRNAs reproduce the ring-like architecture and stall exoribonuclease XRN1 with wild-type-like efficiency. Our top-performing constructs exhibit minimal sequence similarity to known xrRNAs and evade detection by covariance models, yet remain fully functional *in vitro*. Together, our results show that mechanical function can be rationally designed independent of evolutionary ancestry, laying the groundwork for the design of RNA elements that modulate decay and fine-tune the mechanical stability of engineered transcripts.

## 1 Introduction

RNA molecules can encode biological functions through complex three-dimensional topologies that go beyond classical motifs such as stems, hairpins, or junctions. Among these, mechanically active RNAs represent an emerging topic in RNA design: their biological function depends on physically resisting molecular forces rather than on sequence- or structure-dependent recognition or catalysis. To our knowledge, no generalizable rational design framework has been shown to produce such mechanically functional RNA structures.

Exoribonuclease-resistant RNAs (xrRNAs) were initially discovered in the 3’-untranslated region of flaviviruses (genus Orthoflavivirus), where they mediate viral pathogenicity by hijacking and dysregulating the host antiviral response machinery. Upon infection, host 5’ → 3’ exoribonucleases such as XRN1 initiate the degradation of viral RNA but are stalled by the xrRNA element, leading to the accumulation of incomplete degradation products, also known as subgenomic flaviviral RNAs (sfRNAs), which modulate host immune responses. This stalling arises from extreme mechanical anisotropy that acts on the RNA only in the 5’ → 3’ direction [31]. Structural and biophysical studies have shown that this property is encoded by a ring-like fold centered around a three-way junction (3WJ) stabilized by two pseudoknots (PK1 and PK2) and base triplets (Fig. 1, shown here for the single xrRNA of the mosquito-borne Aroa virus (AROAV)). In the three-dimensional structure, the 5’-end is threaded through the RNA core, yielding a topologically closed shape that physically blocks exoribonuclease progression. Since the resulting products do not have a 5’-end protruding from the ring that is long enough for XRN1 to bind to [16], these RNAs are no longer exonuclease substrates and are therefore resistant to further degradation. xrRNAs thus represent a natural example of mechanically active RNA nanostructures in which tertiary topology, as opposed to sequence composition, defines biological function. These properties make xrRNAs a powerful benchmark system for RNA nanomechanics and an especially challenging target for design. The lack of highly conserved xrRNA sequences across viruses leaves no evolutionary blueprint from which to infer design rules. Moreover, the presence of pseudoknots and base triples renders conventional secondary structure design approaches insufficient.

**Figure 1:**
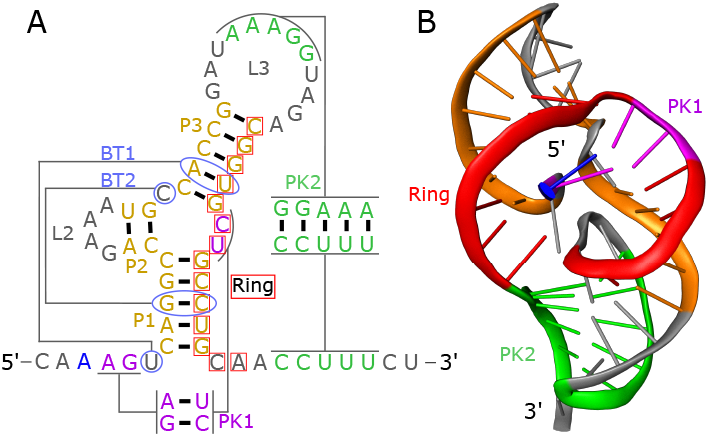
Secondary and tertiary structure of a viral xrRNA. The example shown here represents the single xrRNA element in the 3’UTR of Aroa virus (AROAV) [45]. Corresponding elements in the secondary and tertiary structures are indicated by matching colors. **A**. The secondary structure comprises a three-way junction (3WJ), pseudoknots (PK1, PK2), base triplets (BT1, BT2), and the region forming the ring-like element. **B**. In the corresponding predicted 3D structure, this architectural arrangement collectively establishes the ring topology (red) that encloses the 5’-end (blue) and confers exoribonuclease resistance.

We reasoned that mechanical function could be rationally engineered by directly constraining the topology required for force resistance, rather than relying on sequence conservation. To test this hypothesis, we extracted structural design principles from a pool of representative natural xrRNAs, focusing on the length-correlated features that stabilize the ring-like topology. We then implemented these principles in a mechanics-aware RNA design framework that comprises symbolic structural modeling, adaptive sequence sampling, and molecular dynamics (MD) guided optimization.

Existing RNA design frameworks rely heavily on multiple-sequence alignments (MSAs) and covariance models that capture only the covariation of nested canonical base pairs. Such models, implemented, for example, in `Infernal` [29], are powerful for homology detection but are inherently unable to reproduce pseudoknots, base triples, or other non-nested tertiary contacts that define mechanically active RNAs. Moreover, MSA-driven approaches entangle structural inference with evolutionary conservation, thereby limiting their ability to explore RNA design spaces in which topology, rather than ancestry, defines function.

Conversely, machine learning approaches for RNA design have recently emerged [11]. While such techniques have been proposed for aptamers [12, 46] and ribozymes [22], they depend on large sets of information-rich training sets, which are not available for xrRNAs. As a result, these data-driven methods cannot readily support design strategies based solely on topological or mechanical constraints. Consequently, there is currently no computational framework available that can generate novel xrRNA sequences that are both topologically correct and exonuclease-resistant.

To overcome these limitations, we developed a mechanics-aware RNA design strategy that builds on the `Infrared` framework [47], extending it from classical secondary structure design to include explicit topological and geometric constraints. As a reference for model construction, we used the mosquito-borne Aroa virus (AROAV) xrRNA [45]. Structural features and correlations extracted from natural xrRNAs were encoded as symbolic constraints within the design model. Sequence sampling under these constraints was followed by Monte Carlo optimization toward the target secondary structure, and subsequent *in silico* validation combining 3D modeling and MD-based mechanical testing. Only candidates exhibiting a correctly folded ring-like topology and comparable mechanical stability to the biological reference were selected and *in vitro* transcribed for experimental validation, encompassing exonuclease-catalyzed degradation assays, stop point analysis, and structure probing.

Using this approach, we designed multiple fully synthetic xrRNAs that reproduce the characteristic tertiary topology and stall 5’ → 3’ degradation with efficiencies comparable to those of their biological counterparts. Importantly, our most advanced designs exhibit minimal similarity to known xrRNAs, which is evidenced by evading covariance model detection entirely. Yet, these purposefully designed synthetic xrRNAs remain mechanically robust and fully functional *in vitro*. This demonstrates that the mechanical performance of the molecules can be decoupled from their biological ancestry, establishing a new approach for the *de novo* design of structured RNAs. To our knowledge, this is the first rational strategy to engineer mechanically active RNA devices and moves xrRNAs from a largely descriptive virological curiosity to a tractable design target. The generality of this approach enables the construction of synthetic RNA elements with efficient exonuclease stalling and directional force resistance, allowing for the creation of novel topological architectures not found in nature and with applications in synthetic biology and next-generation RNA therapeutics.

## 2 Material and Methods

### 2.1 *In silico* characterization of biological xrRNAs

To develop an in-depth understanding of biological xrRNAs and construct the constraints used for designs, we first construct a symbolic representation of functional xrRNA. After downloading sequence data of all viruses from the genus Orthoflavivirus from the public National Center for Biotechnology Information (NCBI) GenBank database [6], we selected a representative sequence for each mosquito-borne flavivirus (MBF) species (Tab. S1 and Fig. S1). After aligning the sequences using `mLocARNA` [43, 44], a secondary structure was calculated with `RNAalifold` [2], regions of sequence and structure conservation were extracted, and the total length as well as the lengths of individual structure elements were determined. This information was used to build a symbolic model as shown in Fig. S2A. The model represents the diversity within biological xrRNAs as well as structural correlations. Bases are explicitly included if more than 90% sequence identity is observed in the biological sequences, and all features of the secondary structure are annotated with a range of lengths obtained from the representative sequences. Furthermore, we calculated the correlation between the lengths of the individual structural elements using the Pearson correlation coefficient, shown in Fig. S2B.

### 2.2 Sequence design with Infrared

In the first step, we designed sequences akin to the variety presented in the symbolic xrRNA secondary structure representation shown in Fig. S2. We chose the `Infrared` framework [47] for its flexibility to adapt additional design constraints and objectives. The detailed description of the xrRNA sequence design scheme, following [48], is shown in Fig. S3. Here, we provide a brief overview.

1. **Modeling**. Each position in the symbolic representation can assume the values A, C, G, U, or an empty letter *ε*. The empty letter is analogous to a gap in the aligned sequences, allowing a variation in the length during the design process. A similar idea is implemented in [35]. The designed sequence is then the concatenation of assigned values. Three constraint types induced from the symbolic representation are imposed to restrict the value assignment:
  a. Positions with sequence conservation above 90% are limited to the conserved nucleotide(s).
  b. Nucleotide complementarity is imposed on paired positions in the base structure and in pseudoknots.
  c. To ensure structural integrity, the number of gap characters within components like hairpins is strictly regulated. This ensures that individual element lengths and their mutual correlations remain within the specified ranges (Tables S2 and S3).

Valid assignments are then sampled from a Boltzmann distribution, with sequence weights designed to favor promising xrRNA candidates. The weight is composed of two factors:

a. Structure energy, favoring sequences that exhibit a low folding energy on the target structure. Instead of the full nearest neighbor energy model, we use a base pair energy model previously employed in RNA sequence design [14].
b. The total sequence length is close to the average length of the natural xrRNAs.

Additional constraints are imposed to ensure non-ambiguous sequence sampling.

2. **Monte Carlo Optimization**. To increase the sequence specificity toward the target xrRNA secondary structure, we perform a Monte Carlo post-sampling optimization in two steps: ensemble defect minimization [49] to find sequences that can fold approximately to the target, followed by a maximization of the Boltzmann probability of the sequences folding into the exact target structure. We found that the two-step approach enables faster convergence. Note that the sequence length can change during optimization.

### 2.3 *In silico* selection and validation

The tertiary structures of the candidate xrRNA designs were simulated with `SimRNA` [3], using 25 simulations with 16 replicas and 20 million steps each, guided by secondary structure constraints according to the xrRNA model. After clustering of the 1% lowest energy frames using the scripts provided by `SimRNA`, such that each structure has at most an RMSD of 5 Å to each other structure it is clustered with, the highest occupancy cluster was selected for further analysis. After `QRNAS` [39] minimization, the structure was visually validated with `PyMOL` [37]. Only structures displaying a ring-like topology were considered for further analysis.

As a first test of the structural integrity of the designed xrRNAs, molecular dynamics (MD) simulations were performed for each design and the Root Mean Square Deviation (RMSD) was observed. This was performed using `AMBER24` [4], the RNA.OL3 nucleic acid force field [50] and the TIP3P water model [19]. The simulations were set up using the `Charmm-GUI` solution builder [17, 23, 24] with a 12 Å edge distance and a 0.15 M ion concentration of KCl. The systems were initially minimized over a series of 2500 steps employing the steepest descent method, subsequently followed by an equal number of steps utilizing the conjugate gradient method. Subsequently, the systems were heated to 300 K over 125 ps with a timestep of 1 fs.

All remaining simulations were executed in the NPT ensemble with a temperature of 300 K and a pressure of 1 atm. The system was equilibrated for 125 ps using a 1 fs time step, followed by three replica simulations of 100 ns with a 2 fs time step. The SHAKE algorithm [36] was employed to hydrogen bonds, and long-range electrostatic interactions were calculated using the Particle Mesh Ewald Method [8].

To further test the stability of the ring-like architecture, a procedure to simulate a nanopore-sensing experiment using molecular dynamics was adapted from [31] and [41]. The 5’ end of the structure was placed above a pore, representing the active site of the enzyme, and a force gradient of 1000 pN/μs was applied to the 5’ most nucleotide, pulling it through the pore. Using these simulations, the force required to break up the ring structure can be compared between candidates. The system setup was performed using the implicit solvent builder in `Charmm-GUI` [17, 42]. The parameters were optimized for nucleic acids from the implicit solvent model GBn2 [30], an implicitly modeled salt concentration of 0.15 mol/l, a temperature of 300 K as well as the `AMBER` forcefield OL3 [50] and simulated using the `OpenMM` [9] suite with no cutoff for nonbonded interactions and hydrogen bond constraints for up to 500 ns. `MDAnalysis` [27, 13] was used to determine the unfolding force in post-analysis. It should be noted that the resulting force values highly depend on the simulation parameters, in particular on the slope of the force gradient applied to the 5’ end, and should not be interpreted as a physical force. However, force values can be compared between different structures. To determine a reference force, the procedure was repeated for the natural xrRNA sequence identified in the genome of the Aroa virus, as listed in Table S1. We chose not to select an experimentally determined xrRNA structure to make the comparison to synthetic design candidates as valid as possible. A more detailed description of the setup can be found in Section D of the Supplementary Data.

### 2.4 *In vitro* production of xrRNA candidates

The most promising sequence designs were prepared for validation by *in vitro* XRN1 degradation assays. Using `Infrared`, a unique leader region was designed for each candidate. This sequence had a length of 25 nucleotides and carried at least four unpaired positions at the 5’-end required for XRN1 binding. Furthermore, it contained a single helix with 3 to 5 base pairs to reduce structural interference with the xrRNA fold.

Transcription templates for xrRNA constructs were generated using overlap extension PCR. The resulting products were inserted by restriction-free cloning into plasmid pCR2.1TOPO according to the manufacturer (Thermo Fisher Scientific). The resulting plasmids were amplified in Escherichia coli TOP10 cells, and their sequences were verified by Sanger sequencing. Sequences of individual xrRNA elements are shown in Supplementary Table S4. The transcription units carried a hammerhead ribozyme at the 5’-side and a hepatitis delta virus ribozyme at the 3’-side to ensure homogeneous end generation after transcription [28]. To generate a template for *in vitro* transcription, the desired plasmid region containing a T7 promoter and the ribozyme cassette including the xrRNA sequence was amplified by PCR. Alternatively, the plasmid was linearized with EcoRV and used as a template. *In vitro* transcription was performed with T7 polymerase, and the resulting RNA was purified via denaturing 10% polyacrylamide gel electrophoresis (PAGE) and subsequent elution in 200 mM NaCl, 10 mM Tris (pH 7.5), 1 mM EDTA, followed by ethanol precipitation.

### 2.5 Degradation resistance assay

Cleavage of the hammerhead ribozyme results in an RNA molecule carrying a 5’OH group. Since XRN1 only recognizes RNA with a 5’-phosphate group, the *in vitro* transcribed RNA was phosphorylated at the 5’-end using the T4 polynucleotide kinase (3’-phosphatase minus) (New England Biolabs). After phenol-chloroform extraction and ethanol precipitation, the RNA was refolded in a temperature gradient from 90 °C to 37 °C to ensure proper folding of the xrRNA structure. 500 ng of the refolded RNA was incubated with 1U XRN1 (New England Biolabs) in the corresponding NEBuffer™ 3 for 1h at 37 °C. As a negative control, the reaction was incubated in the absence of XRN1. The reaction was stopped by the addition of 3x RNA loading dye (10 mM Tris-HCl (pH 7.6), 80 % (v/v) formamide, 0.25 % (w/v) bromophenol blue, 0.25 % (w/v) xylene cyanol) and 15 mM EDTA. Reaction products were separated on a 10 % denaturing polyacrylamide gel and visualized by Sybr™Gold (Thermo Fisher Scientific) staining for 30-60 min according to the manufacturer’s specifications. An alkaline hydrolysis ladder of each xrRNA candidate was prepared by incubation for 5 min at 95 °C in the presence of 50 mM Na_2_CO_3_/NaHCO_3_ and included as a size standard.

### 2.6 Quantification of XRN1 resistance

The intensity of the gel bands representing the substrate and reaction products (XRN1-resistant transcript part) was determined using ImageQuant TL (GE Healthcare). Since the yield of the RNA substrate 5’-phosphorylation is not 100%, the intensity of the band of non-phosphorylated RNA was subtracted from the untreated RNA intensity before calculating the ratio of substrate to XRN1-resistant RNA. The obtained ratio of the wildtype xrRNA was set to 100%, and the ratios of the individual constructs were normalized to it.

### 2.7 Structure analysis by in-line probing

*In vitro* transcribed RNA was radioactively labeled at the 5’-end using *γ*-^32^P-ATP (Hartmann Analytic), 8 mM DTT, 1x PNK buffer, and T4 Polynucleotide kinase (New England Biolabs). The reaction product was purified via electrophoresis on a 10% denaturing polyacrylamide gel. 10 pmol RNA (30,000 cpm) were treated according to the refolding protocol described above. Inline probing buffer (50 mM Tris/HCl (pH 8.5), 20 mM MgCl_2_, 100 mM KCl) was added. In-line probing was performed for 24 h at 37 °C as described [33, 20, 10], including negative control and size standards (generated by RNase T1-catalyzed G-specific cleavage and partial alkaline hydrolysis). Reactions were stopped by the addition of 3x colorless RNA loading dye, including 13.5 mM EDTA. Reaction products were separated on a 10% polyacrylamide gel and visualized with a Typhoon FLA 9410 PhosphorImager (Cytiva).

### 2.8 XRN1 stop point analysis

XRN1 reaction was stopped by heat inactivation at 70 °C for 10 min. The exonuclease-resistant RNA was dephosphorylated with Antarctic phosphatase (New England Biolabs) according to the manufacturer’s protocol, except for a prolonged incubation time of 1 h. The RNA was purified using the Monarch ^®^ RNA cleanup kit (New England Biolabs) and eluted in 6 μl sterile and RNase-free water. The RNA was used for 5’-adapter ligation with RtcB ligase according to the Led-Seq protocol [21]. The transcripts were PCR-amplified to introduce adapter sequences for Amplicon sequencing (Azenta). PCR products were purified on a 10% agarose gel, eluted with GeneJET gel extraction kit (Thermo Scientific) and analyzed by Amplicon sequencing (Azenta). This sequencing resulted in read numbers of up to 52.000 for the individual resistant RNA designs.

Sequences were extracted from the raw reads and mapped to the complete xrRNA sequences using `Segemehl` [15]. For all analyzed constructs, distinct and unambiguous stop point positions were identified as indicated in Fig. 2, 4, and 5 with detailed results shown in Fig. S4.

**Figure 2:**
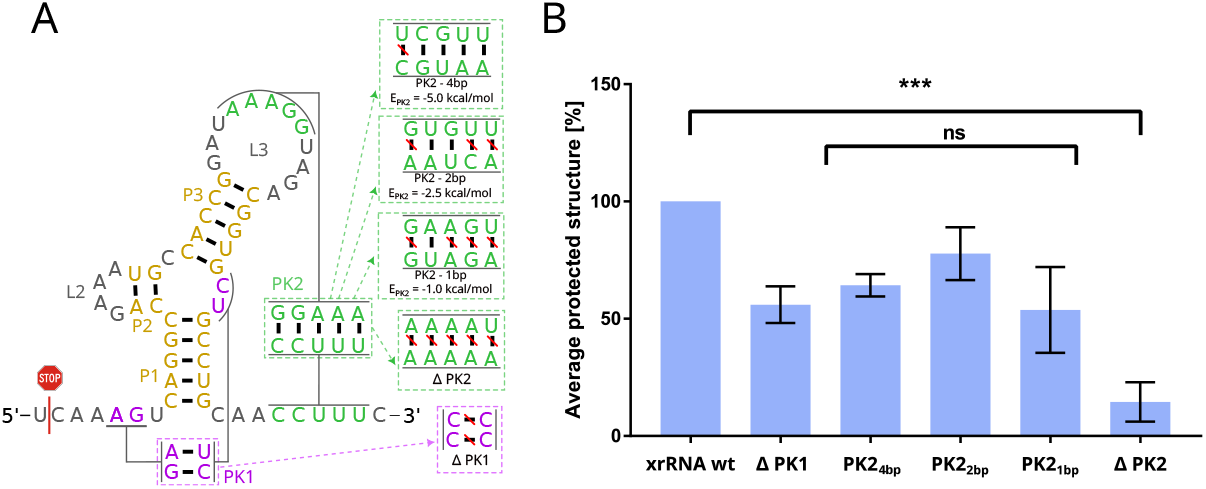
XRN1 degradation assay quantifying the mechanical contribution of pseudoknots to AROAV xrRNA function. The wild-type sequence and structure were systematically modified to either delete or progressively weaken PK2, or to prevent PK1 formation. The identified XRN1 stop point of the wild-type xrRNA is indicated by a red stop symbol. Secondary structures and corresponding sequences of the mutant constructs (panel A) are shown alongside their relative protection against XRN1 degradation (panel B). Statistical significance was assessed using Welch’s t-test and one-way ANOVA (P *<* 0.05).

**Figure 3:**
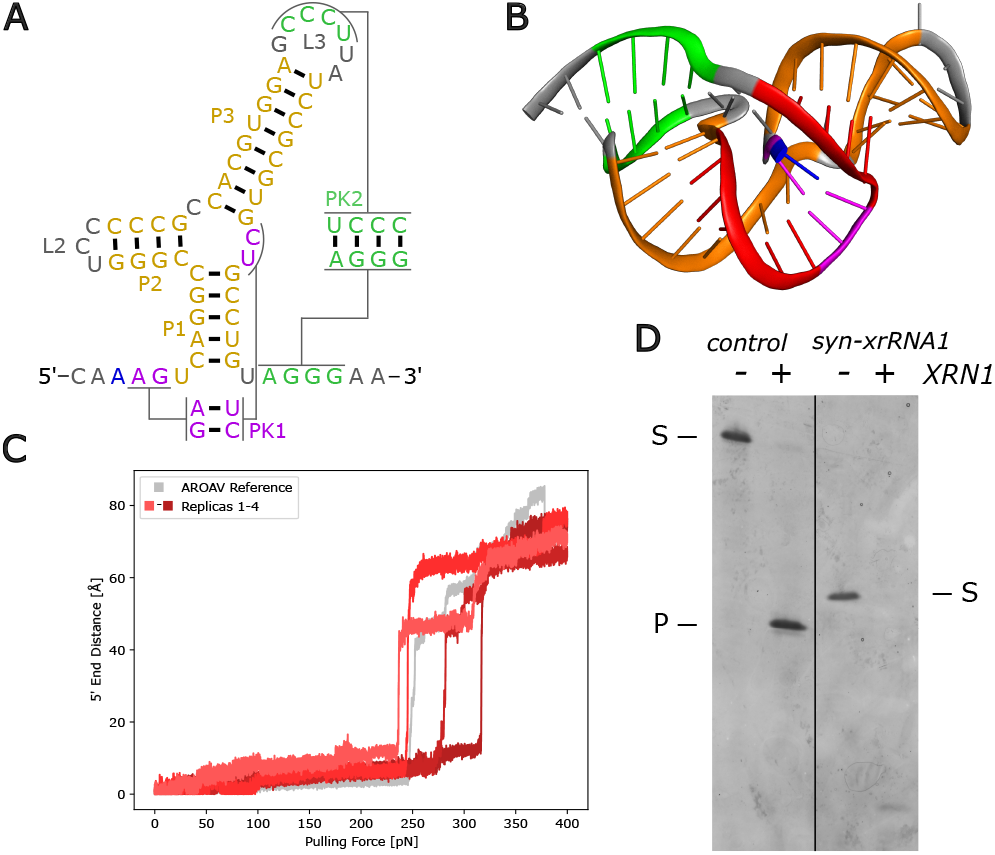
Structure prediction and experimental validation of syn-xrRNA1. **A**. Secondary structure model according to the base pairs encoded in the `Infrared` model. The 3-way junction is flanked by three GC-rich helical domains (P1 – P3, yellow). Pseudoknot 1 (PK1) is labeled in purple, pseudoknot 2 (PK2) in green, with the 5’ end in blue. **B**. 3D structure predicted by `SimRNA` using a coloring scheme similar to the one used in panel A. The ring-like structure (red) with the protruding 5’-end of the RNA (blue) is formed, representing the expected structural organization. **C**. Computational pulling-force experiment mimicking XRN1 activity. Four replicas (red) are shown for syn-xrRNA1. The wild-type AROAV xrRNA structure model was used as a reference (gray). The data indicate a comparable resistance of the designed and wild-type xrRNA elements. **D**. XRN1 *in vitro* degradation assay. While the AROAV xrRNA transcript showed a strong resistance against the exonuclease, syn-xrRNA1 was efficiently degraded, indicating that the construct does not represent a functional exonuclease-resistant RNA. We note that different xrRNAs vary in length, hence travel different distances in the PAGE. S, substrate transcript (AROAV: 102 nts; syn-xrRNA1: 84 nts); P, product of XRN1 degradation (AROAV: 79 nts).

**Figure 4:**
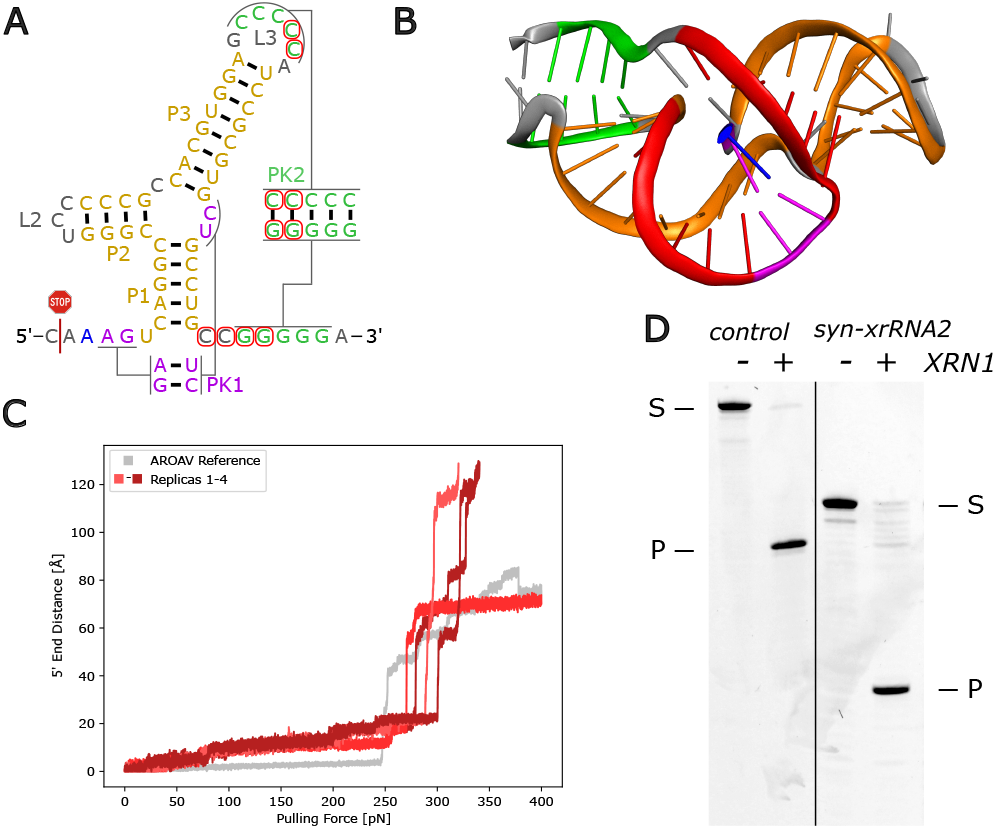
Design and validation of syn-xrRNA2. **A**. Secondary structure model encoded in the Infrared design framework. Nucleotide changes relative to syn-xrRNA1 are highlighted by red circles. In pseudo-knot 2 (PK2), one additional C-G base pair was introduced and one U-A pair was replaced by a G-C pair, resulting in a substantial stabilization of this interaction. The experimentally determined XRN1 stop position, obtained by amplicon sequencing, is indicated by a red stop symbol. **B**. `SimRNA`-based prediction of the 3D structure, showing the formation of the ring-like topology (red) with the protruding 5’-end (blue). **C**. *In silico* simulation of the structural resistance to the pulling force exerted by XRN1. As in Fig. 3, four independent replicas were performed and compared with the AROAV xrRNA. **D**. Exonucleolytic activity of XRN1 on *in vitro* transcribed syn-xrRNA2. The construct displays exonuclease resistance comparable to that of the AROAV xrRNA control, indicating that the introduced changes contribute to stabilization of the transcript. S, substrate RNA (AROAV: 102 nts; syn-xrRNA2: 85 nts), P, product RNA (AROAV: 79 nts; syn-xrRNA2: 61 nts).

**Figure 5:**
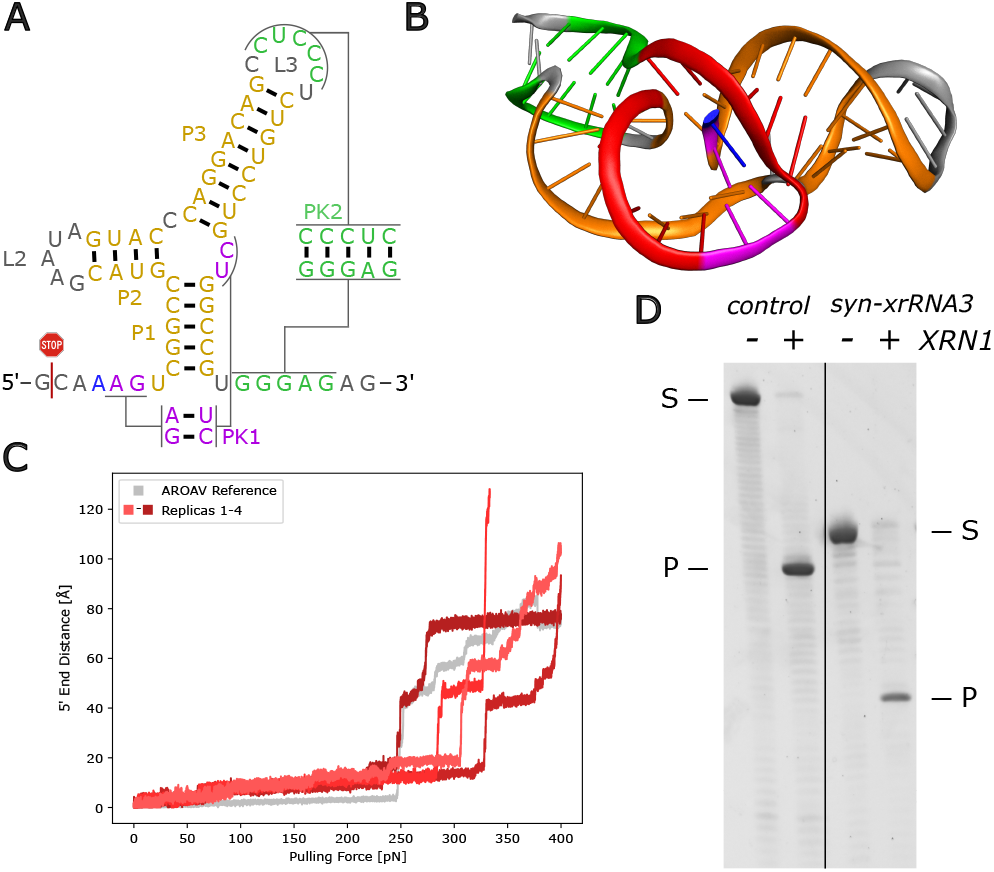
Design and characterization of syn-xrRNA3. **A**. Secondary structure representation defined by the base pairs encoded in the Infrared design model. The XRN1 stall position, determined by amplicon sequencing, is indicated by a red stop symbol. **B**. `SimRNA`-based prediction of the 3D structure, shown using a color scheme consistent with panel A. The model adopts the expected ring-like topology with the 5’-end protruding from the fold. **C**. *In silico* simulations probing the resistance of the ring-like structure to an applied pulling force, compared with the AROAV reference (4 replicas). The construct exhibits strong mechanical robustness with a mean unfolding force of 295.5 pN. **D**. *In vitro* XRN1 degradation assay, with the AROAV xrRNA used as a positive control. XRN1-mediated degradation yields a stable reaction product (P), indicating exonuclease resistance of syn-xrRNA3. S, substrate transcripts (AROAV: 102 nts; syn-xrRNA3: 83 nts), P, resistant reaction products (AROAV: 79 nts; syn-xrRNA3: 63 nts).

### 2.9 Software availability

All scripts developed for this study are available on GitHub at https://github.com/ViennaRNA/xrRNA-design-code.

## 3 Results

### 3.1 Pseudoknot architecture encodes force resistance

The hallmark of xrRNA function is its ability to withstand 5’ → 3’ exonucleolytic degradation through mechanical resistance encoded in a topologically closed RNA fold. Previous structural and biochemical studies have shown that pseudoknots are indispensable for such a fold. They lock the 5’-end within the ring-like topology that confers mechanical resistance [32], suggesting some concerted or cooperative effect of the PKs.

To dissect how individual pseudoknots contribute to this property, we systematically perturbed the pseudoknot architecture of the Aroa virus (AROAV) xrRNA, which we used as a reference system for mechanical benchmarking. Five mutant constructs were generated in which either the first pseudoknot (PK1) or the second (PK2) was deleted, or, in the case of PK2, progressively weakened by removing base pairs (Fig. 2). All constructs contained a designed leader sequence susceptible to XRN1 degradation, allowing direct quantification on denaturing PAGE. Degradation assays are shown in Fig. S5.

Progressive weakening of PK2 reduced resistance to 50–75 % of wild-type levels as long as at least one base pair remained intact. Complete deletion of PK2, however, abolished resistance entirely, whereas deletion of PK1 resulted in a partial stalling effect. This identifies PK2 as the primary determinant of mechanical resistance in the AROAV xrRNA. Its presence enables closure of the ring-like topology that physically traps the 5’-end, while PK1 and the associated base triplet interactions contribute additional stabilization, resulting in an additive effect on exonuclease resistance. Notably, the total free energy of the pseudoknot interactions was less predictive of resistance than their specific geometric arrangement, indicating that topology, rather than thermodynamic strength alone, governs mechanical function. These experiments consistently demonstrate that the mechanical integrity of xrRNAs is encoded in a hierarchical pseudoknot architecture. PK2 defines topological closure, while PK1 supports structural integrity. We used these insights to derive the design rules that guided the rational engineering of artificial xrRNAs described below.

### 3.2 Rational design of artificial xrRNAs with wild-type-like mechanical function

Building on our observations from the pseudoknot perturbation analysis, we constructed a symbolic model of MBF xrRNAs that captures the essential sequence and structural features of natural, endogenous xrRNAs (Fig. S2A). The model includes the characteristic three-way junction (3WJ), encompassed by three stems (P1-P3), and two pseudoknots (PK1 and PK2), which are required for the formation of the ring-like fold. Bases were maintained at positions with *>* 90% sequence identity. We also encoded quantitative correlations between stem and loop lengths. Notably, strong negative correlations between the length of P3 and loop L3 suggest that natural xrRNAs display a balanced geometry between the long stem and its capping loop. This is likely a constraint for preserving the ring topology. These correlations were included in the design model as quantitative length relationships.

Our analysis of natural xrRNAs revealed substantial sequence variation and divergence in the length of structural elements, although some regions exhibit high sequence conservation. The region around the central 3WJ is highly conserved, comprising exclusively C-G pairs within the junction. In contrast, P2 and L2 exhibit high diversity in MBF xrRNAs, with some sequences lacking P2 completely. A similar length variation was also identified for PK2, consisting of 3 to 8 base pairs. Conversely, P1 is fully conserved in terms of length and base composition.

We used these insights to construct a model for sequence sampling and optimization using the `Infrared` design framework, which allows the inclusion of structural and energetic constraints during sequence generation. The first construct, syn-xrRNA1, was designed conservatively to retain most of the MBF xrRNA consensus while allowing for minor sequence variation. *In silico* folding and three-dimensional modeling predicted a well-formed ring topology (Fig. 3 A and B) that remained stable over extended molecular-dynamics (MD) simulations (Fig. S6). We used steered molecular dynamics simulations, mimicking the pulling action of XRN1, to compare the stability of syn-xrRNA1 to the natural AROAV reference. Using our setup, we measured an unfolding force of 263.5 pN, which closely matches the 250.0 pN determined for the reference. While these values are highly dependent on the simulation parameters and should not be interpreted as physical forces, the comparison indicates a mechanical performance similar to natural xrRNAs(Fig. 3 C).

Structural evaluation by in-line probing confirmed that syn-xrRNA1 forms the expected structure but exhibits incomplete pseudoknot formation (Fig. S7). In accordance with this structural weakness, XRN1 degradation assays showed reduced resistance compared to the wild-type control (Fig. 3 D). This confirmed that a correct topology was achieved, although the local reinforcement of PK2 remained suboptimal.

Building on these results, we increased the stability of PK2 by replacing the single U-A pair by C-G and the addition of an extra C-G pair. Furthermore, we extended the linker between P1 and PK2 to improve the geometry of the ring-like structure. The resulting design, syn-xrRNA2 (Fig. 4 A and B), displayed improved mechanical stability in steered MD simulations, requiring 284.6 pN to disrupt the ring-like fold (Fig. 4 C). Interestingly, this value surpasses the stability of the wild-type reference. Improved stability was also observed *in vitro*, where the band in the degradation assay representing the protected RNA showed an intensity similar to the control (Fig. 4 D). In addition, some degradation intermediates are visible in the corresponding gel lane. Moreover, inline probing verified full PK2 formation, although PK1 still showed some fraying (Fig. S7). The improvements made to the design were also added to the constraints model for all further xrRNA designs.

To further characterize this xrRNA construct, the 5’-end of the resulting stable degradation product was determined, representing the stop point of the XRN1-catalyzed degradation. The position is indicated in Fig. 4 A by a red stop symbol and is located two positions upstream of PK1, corresponding to the previously reported stop points of natural MBF xrRNA elements [18]. These results show that the design rules derived from natural xrRNAs are sufficient to reproduce wild-type-like mechanical performance. The successful generation and validation of syn-xrRNA1 and syn-xrRNA2 establish a direct, rational link between pseudoknot geometry, topological closure, and mechanical function. This enables the *de novo* design of mechanically active RNA structures beyond natural sequence constraints.

### 3.3 *De novo* engineering independent of sequence ancestry

To explore the full potential of the design approach outlined above, we asked whether mechanical function can still be maintained when all evolutionary sequence information is removed. To this end, we set up an MBF xrRNA model that minimizes sequence constraints during sampling. While the structural architecture, i.e., the 3WJ, stems P1-P3, the pseudoknots and their corresponding length relationships were preserved, only the nucleotides that form PK1 and the base triplets remained fully constrained. The base pairs forming the central 3WJ were allowed to vary, constrained only to favor G-C pairs to maintain the stability of the fold. In addition, empirical feedback from the first round of designs was incorporated to allow for efficient ring closure. Specifically, we assigned higher weights to pseudoknot formation and adjusted the linker geometry between P1 and PK2.

This new model produced sequences with no detectable similarity to known xrRNA elements when analyzed with `Infernal` covariance models or other state-of-the-art RNA homology search methods [26, 7]. Thus, the resulting candidates qualify as *de novo* xrRNAs that share topological and mechanical, but not sequence, ancestry with xrRNAs found in nature.

Among the resulting design candidates, we selected construct syn-xrRNA3 for detailed evaluation (Fig. 5). *In silico* folding and 3D modeling confirmed a correctly threaded ring topology consistent with that of MBF xrRNAs (Fig. 5 A, B). Steered molecular-dynamics simulations revealed an unfolding force of approximately 295.5 pN, exceeding both syn-xrRNA2 and the wild-type reference (Fig. 5 C), suggesting that mechanical robustness is preserved or even enhanced, despite the absence of recognizable sequence motifs. In accordance with these calculations, the construct exhibited a strong resistance against XRN1-mediated degradation, resulting in a resistant reaction product (Fig. 5 D). The corresponding stop point was characterized as described and is indicated in Fig. 5 A. As observed for syn-xrRNA2, some minor degradation intermediates are also visible for syn-xrRNA3.

Comparison of covariance model scores across all designs highlights the fundamental novelty of syn-xrRNA3 (Fig. 6). CM scores of the artificial designs are compared to randomly sampled sequences that fulfill only the base pair complementarity constraint. While syn-xrRNA1 and syn-xrRNA2 score within the range of natural xrRNAs, syn-xrRNA3 falls within the distribution of random sequences, i.e., it only satisfies structural constraints. Its slightly higher-than-average covariance score can be explained by the tendency of secondary-structure–optimized sequences to contain a greater proportion of C–G base pairs and adenosines at unpaired positions, which is also observed in natural xrRNAs. A detailed analysis of the proximity to biological xrRNAs using covariance models is shown in the Supplementary Material (Fig. S8 and Tab. S5).

**Figure 6:**
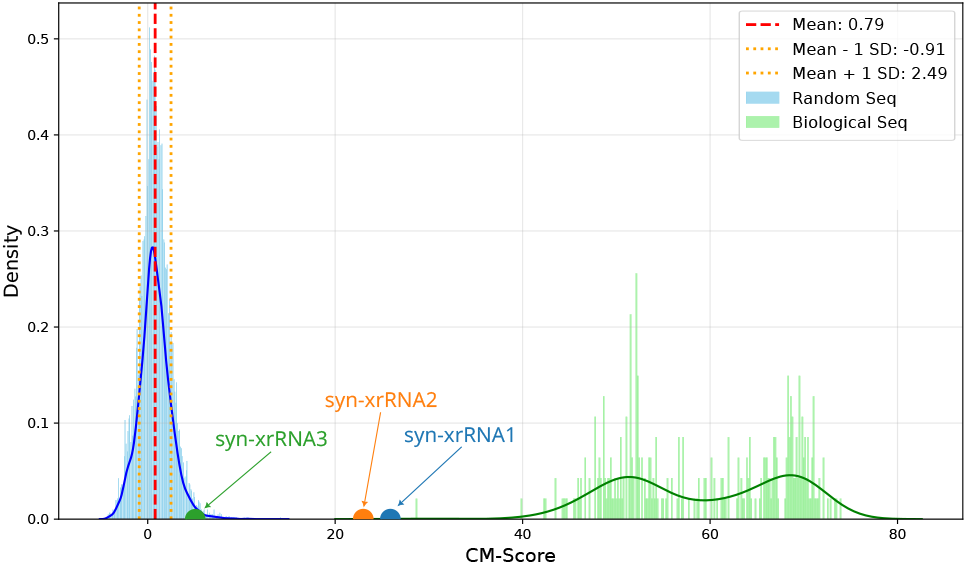
Distribution of covariance model scores for random sequences and naturally occurring xrRNAs. CM scores were calculated by evaluating random sequences that satisfy the structural constraints used in the design process (blue distribution, left), and naturally occurring viral xrRNA sequences (green distribution, right) against a covariance model trained on MBF xrRNA sequences. The scores of the synthetic design constructs syn-xrRNA1 (blue), syn-xrRNA2 (orange) and syn-xrRNA3 (green) are depicted as dots on the x-axis, indicating that they evade detection by the MBF xrRNA CM.

To the best of our knowledge, this establishes syn-xrRNA3 as the first fully artificial, functional xrRNA. The construct operates through the same topological mechanism as natural xrRNAs, despite being derived entirely from rational design. These results show that mechanical performance can be decoupled from biological ancestry and that topology-encoded resistance is a tunable property in RNA design. Notably, our data demonstrate that force resistance is accessible through *de novo* design based solely on structural and energetic constraints.

## 4 Discussion

In this work, we present a rational design strategy for mechanically active RNAs and demonstrate its potential by engineering synthetic xrRNAs that match the performance of their natural counterparts. By systematically perturbing the pseudoknot architecture of a model AROAV xrRNA, we identified structural features and derived design rules that encode resistance to 5’ → 3’ exonucleolytic degradation through topological closure rather than sequence-specific interactions. These rules were then implemented in a mechanics-aware design framework that combines symbolic structural modeling, length-correlation constraints, and molecular dynamics-based mechanical selection. Using this approach, we generated synthetic xrRNAs that reproduce the defining mechanical properties of their natural counterparts and, critically, a fully *de novo* construct that remains functional despite lacking detectable evolutionary ancestry. The results demonstrate that RNA mechanical resistance can be specified directly at the level of topology and geometry, independent of sequence conservation.

The xrRNA pseudoknot perturbation experiments complement and extend earlier observations from other viruses [40], supporting a hierarchical organization that ultimately determines ring closure. PK2 manifested as a critical gatekeeper of mechanical resistance. Gradual weakening of PK2 had only moderate effects, whereas complete deletion abolished stalling of the 5’ → 3’-exonuclease, resulting in complete degradation. In contrast, deletion of PK1 reduced, but did not eliminate, XRN1 resistance in the AROAV xrRNA transcript, suggesting that PK2 and the surrounding base triples can partly compensate for the loss of PK1. This aligns with earlier comparative studies showing that different flaviviral xrRNAs distribute functional load differently among PK1, PK2, and associated tertiary contacts [5, 1, 25]. The perturbation experiments support a model in which overall architecture, rather than any single motif, determines xrRNA resistance.

Building on these mechanistic insights, we developed a symbolic model of MBFV xrRNAs that incorporates not only secondary structure but also element-wise length distributions and correlations. This representation allowed us to treat xrRNA design as a constrained search over topological architectures, rather than as a sequence-optimization problem guided by multiple sequence alignments. Integrating the `Infrared` method for sequence sampling with ensemble-defect minimization and molecular dynamics-based force profiling yielded a closed design-validation loop in which only candidates predicted to have plausible topology and high mechanical robustness were selected for experimental validation. In practice, this approach focuses expensive experimental effort on the most promising designs.

A key implication of this study is that mechanical function can be decoupled from evolutionary sequence ancestry. While syn-xrRNA3 was designed under minimal sequence constraints, it scores like a random sequence under covariance models trained on known xrRNAs (Fig. 6), and yet displays robust XRN1 stalling and simulated unfolding forces that exceed the wild-type reference (Fig. 5). This finding demonstrates that topology and mechanical design principles are sufficient to specify function, and that traditional homology-based approaches will systematically miss large regions of functional RNA design space [38, 34].

While the first applied design syn-xrRNA1 did not yet result in a structure sufficiently resistant against XRN1, the enhanced PK2 and the improved ring geometry of syn-xrRNA2 led to an exonuclease-resistant shape similar to the wild-type AROAV element. The exonuclease stop point in syn-xrRNA2 is shifted one position downstream compared to the natural element (Fig. 4), suggesting that the 3D structure of this RNA element is less rigid than that of the AROAV xrRNA, allowing for more ring flexibility. In syn-xrRNA3, in contrast, the stop position is identical to the control xrRNA, indicating that this design has a stability comparable to the natural structure (Fig. 5). This observation underlines the functionality of our novel structural design approach.

In this work, we demonstrate that functional RNA structures can be designed rationally, without relying on large multiple sequence alignments or high-throughput screening, provided that the relevant mechanical and topological constraints are correctly specified and encoded. While we focused here on xrRNAs as a benchmark for force-dependent function, the same strategy can, in principle, be applied to other topologically defined RNA elements, such as G-quadruplexes and kink-turns. Applying similar techniques to other RNA families would enable a more systematic exploration of how mechanically active RNAs can be used to modulate degradation or processing pathways.

Finally, the ability to engineer *de novo* xrRNAs has direct implications for RNA-based therapeutics and synthetic biology. Natural xrRNAs and viral 3’-UTR fragments have already been used to modulate RNA stability [5], but their utility is constrained by native sequence context. The approach presented here enables the construction of tailor-made, mechanically defined RNA elements that can be embedded into diverse RNA contexts without relying on native viral sequences. Such mechanically defined RNA modules may be used to enhance transcript stability or protect specific RNA regions from exonucleolytic decay. In this sense, the present study offers a starting point for the rational incorporation of mechanically functional RNA motifs into synthetic and therapeutic RNA constructs.

## Author contributions statement

LS and KG developed `Infrared` models for synthetic xrRNAs design, performed 3D predictions, and compared designed and natural xrRNA sequences. DS curated representative xrRNA sequences and analyzed sequence conservation and local secondary structure features. DS and LS designed AROAV mutants with different pseudoknot strengths. LS performed molecular dynamics simulations to assess xrRNA stability. JW performed *in vitro* XRN1 degradation and in-line probing assays and conducted stop-point analysis experiments, which LS computationally evaluated. TK initially tested the *in vitro* experimental protocols and performed preliminary experiments. HTY and ILH provided input on the `Infrared` RNA design process. HTY participated in guiding the *in silico* design. MM and MTW conceived the study, supervised the work, and secured funding. LS, JW, and HTY drafted the manuscript. MM and MTW edited and revised the manuscript. All authors approved the final manuscript.

## Acknowledgments

We thank Niklas von Malottki and Anna Gerbsch for their assistance in the *in vitro* analysis of XRN1-mediated degradation. This work was supported by the Deutsche Forschungsgemeinschaft (DFG) [project Mo 634/24–1 to MM], the Austrian Science Fund FWF [project I 6440-N to MTW], and the Asean-European Academic University Network (ASEA-UNINET). We acknowledge support from the Deutsche Forschungsgemeinschaft (DFG) and Leipzig University within the program of Open Access Publishing.

