## Supplementary Data for "Rational design of mechanically active RNAs: de novo engineering of functional exoribonuclease-resistant RNAs"



### A Reference Sequences

Parameters for the symbolic models were derived from natural xrRNA sequences in mosquito-borne flaviviruses (MBFV, genus *Orthoflavivirus*). Viral genomes and corresponding annotations were retrieved from the public NCBI Genbank database [3], and xrRNA loci were extracted with `Infernal` [8]. For each of the viruses listed in Tab. S1, the corresponding RefSeq isolate was selected as a representative. If no RefSeq isolate was available, the isolate with the longest annotated 3' UTR was chosen.

For model parametrization, the first xrRNA occurrence (xrRNA1) from each representative genome [13] was used to compute the multiple sequence alignment shown in Fig. S1 with `mLocARNA` [12].

**Table S1:** Representative MBFVs used to build the xrRNA models, and relative position of the first xrRNA in the viral 3' UTR.

| Virus Name (Abbreviation) | Accession Number | Position in 3' UTR |
|---|---|---|
| Bagaza virus (BAGV) | MF380429.1 | 117-180 |
| Aroa virus (AROAV) | NC_009026.2 | 29-85 |
| Banzi virus (BANV) | AY326407.1 | 151-207 |
| Dengue virus type 1 (DENV1) | NC_001477.1 | 123-175 |
| Dengue virus type 2 (DENV2) | MT899084.1 | 30-91 |
| Dengue virus type 3 (DENV3) | MZ284953.1 | 50-111 |
| Dengue virus type 4 (DENV4) | MK858146.2 | 32-90 |
| Japanese encephalitis virus (JEV) | ON875960.1 | 49-110 |
| Kokobera virus (KOKV) | OL347997.1 | 34-88 |
| Koutango virus (KOUV) | OQ067500.1 | 67-128 |
| Kunjin virus (KUNV) | KX394396.1 | 99-161 |
| Murray Valley encephalitis virus (MVEV) | KF751869.1 | 91-154 |
| New Mapoon virus (NMV) | NC_032088.1 | 37-90 |
| St. Louis encephalitis virus (SLEV) | JQ957869.1 | 36-97 |
| Tembusu virus (TMUV) | OQ473625.1 | 98-156 |
| Uganda S virus (UGSV) | AY326410.1 | 47-104 |
| Usutu virus (USUV) | JQ219843.1 | 143-204 |
| Wesselsbron virus (WESSV) | EU707555.1 | 160-222 |
| West Nile virus (WNV) | MZ605382.4 | 126-188 |
| Yellow fever virus (YFV) | MW158349.1 | 111-181 |
| Zika virus (ZIKV) | MH675626.1 | 16-74 |

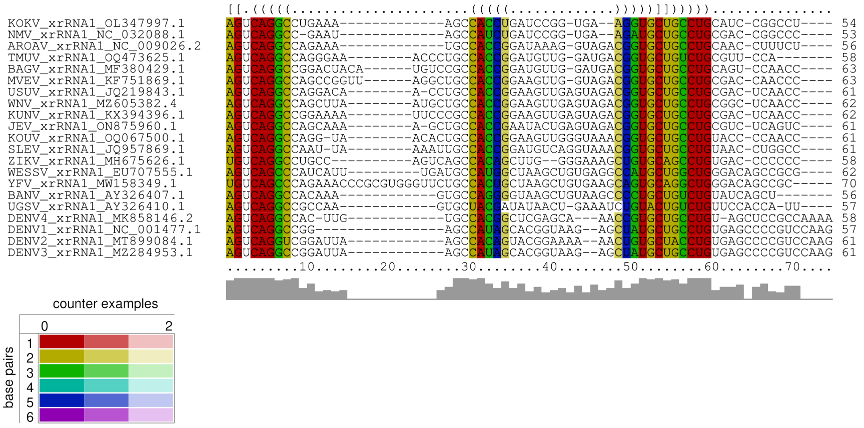


**Figure S1: Structural multiple sequence alignment of natural xrRNAs.** Structural alignment and consensus secondary structure of natural xrRNA sequences from representative MBFV isolates calculated with `mLocARNA`. Square brackets depict pseudoknot interactions of PK1. PK2 is not predicted in this consensus structure.

### B Correlation between xrRNA structural elements

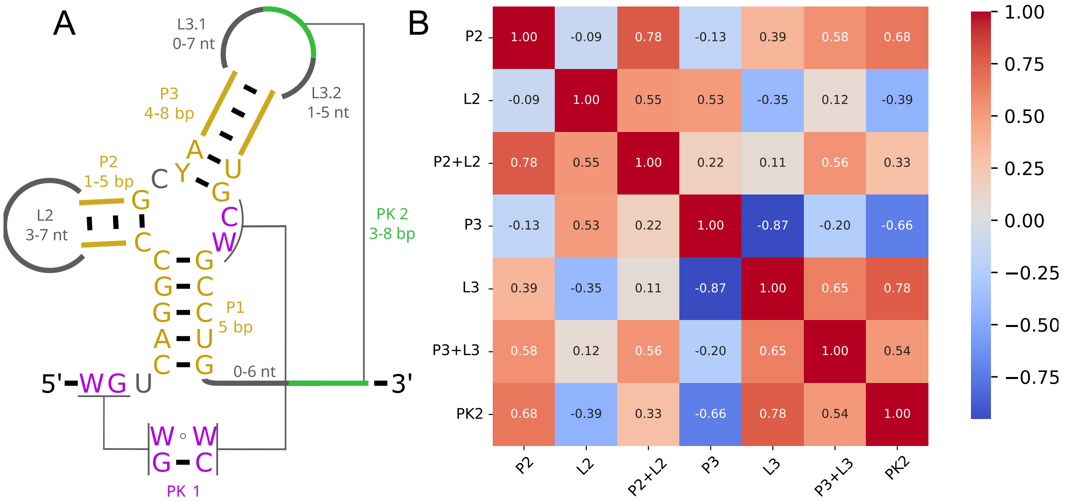


**Figure S2: Length correlations between xrRNA structural elements.** A. Symbolic representation of the structural diversity of MBFV xrRNAs. Conserved elements are highlighted in ochre (stems P1-P3), grey (loops L2 and L3), and purple/green (Pseudoknots PK1 and PK2). Positions with >90 % sequence conservation are shown using IUPAC nucleotide symbols. B. Pearson correlation coefficients quantifying the association between the lengths of conserved structural elements in the selected MBFV xrRNA sequences.

### C xrRNA design with `Infrared`

Conventional RNA secondary structure design optimizes sequence compatibility with a defined target fold, typically by performing a stochastic walk in sequence space starting from an initial seed sequence, as reviewed in [2]. Common objective terms include the distance between the predicted minimum free-energy (MFE) structure and the target, the Boltzmann probability of the target structure, and ensemble-defect measures.

In our xrRNA setting, an additional constraint is that the search must remain within the sequence space defined by the symbolic representation. To enforce this throughout the design

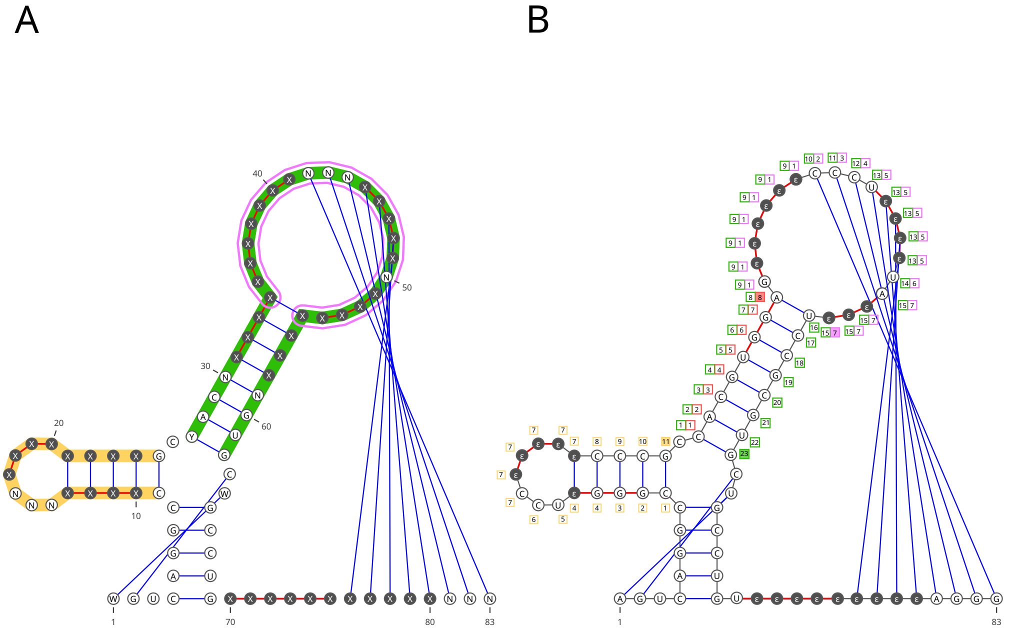


**Figure S3: Visualization of the `Infrared` model.** A. Variables used in the design model. Position-specific domains are indicated using IUPAC nucleotide codes; the additional symbol $X$ denotes the extended domain $\{A, C, G, U, \epsilon\}$. A red backbone marks regions subject to a unique sequence constraint, whereas colored outlines indicate length-correlation constraints between regions. B. Variable assignment for the MBFV model corresponding to the design syn-xrRNA1. Auxiliary variables used to count region lengths are shown adjacent to the base positions as color-coded squares. The terminal variable representing the total length of each region is highlighted as a filled square.

procedure, we adopted the `Infrared` framework [14], which formulates sequence generation as a weighted constraint satisfaction problem (CSP) specified by $(\mathcal{X}, \mathcal{D}, \mathcal{C}, \mathcal{F})$. Here, each variable $X_i \in \mathcal{X}$ corresponds to a sequence position and takes values from a domain $D_i \in \mathcal{D}$, typically $\{\mathtt{A}, \mathtt{C}, \mathtt{G}, \mathtt{U}\}$. A valid assignment (sequence) $x = (X_i \mapsto x_i)_{i \in |\mathcal{X}|}$, where $x_i \in D_i$, is a mapping of variables to values satisfying all constraints $\mathcal{C}$. In addition, each valid assignment is evaluated by the linear contribution of features $\mathcal{F}$, $E(x, \alpha) = \sum_{F \in \mathcal{F}} \alpha_F F(x)$ where $\alpha_F$ is the weight of feature $F$. `Infrared` provides an efficient solution to sample a valid assignment $x$ from the Boltzmann distribution with the probability $\mathbb{P}(x) \propto \exp(E(x, \alpha))$.

We used `Infrared` to design variable-length candidate sequences that fold into a pseudoknotted target structure. Because most energy models do not support pseudoknots, the full fold was decomposed into three noncrossing structures $S$, $S_{PK_1}$, $S_{PK_2}$ of length $n$, which denotes the maximum allowed sequence length.

**Base model construction.** Let $n$ denote the maximum xrRNA length implied by the symbolic representation (by considering the maximum possible size of each region). We first consider the variable set $\mathcal{X}_{\text{xrRNA}} = \{X_1, \ldots, X_n\}$ representing $n$ positions of xrRNA ($n = 83$) with domain set $\mathcal{D}_{\text{xrRNA}} = \{D_1, \ldots, D_n\}$. Each variable $X_i$ can take a value from the domain $D_i = \{\mathtt{A}, \mathtt{C}, \mathtt{G}, \mathtt{U}, \epsilon\}$ where $\epsilon$ is the empty character used to represent different length sequences. We limit the domain of certain variables to $\{A, C, G, U\}$, *i.e.* `N` in Fig. S3, to ensure that the minimum size of each region is achieved when the rest of variables have a value of $\epsilon$. As an example, the hairpin L2 has a size ranging from 3 to 7 nt. We expressed in `Infrared` with 7 variables $\{X_{14}, \ldots, X_{20}\}$ with the letter $\epsilon$ excluded from the domain of the variables $X_{14}$, $X_{15}$, and $X_{16}$.

**Base pair complementary.** Canonical pairing was enforced for all base pairs in the xrRNA secondary structure $S$ as well as in the pseudoknot components $S_{\text{PK1}}$ and $S_{\text{PK2}}$. Accordingly, admissible base pairs were restricted to

$$\mathcal{BP} = \{(\mathtt{A}, \mathtt{U}), (\mathtt{C}, \mathtt{G}), (\mathtt{G}, \mathtt{C}), (\mathtt{G}, \mathtt{U}), (\mathtt{U}, \mathtt{A}), (\mathtt{U}, \mathtt{G}), (\epsilon, \epsilon)\}. \quad (1)$$

This requires imposing a base pair complementary constraint (blue edge in Fig. S3) on each base pair

$$\mathcal{C}_{\text{comp}} = \{\text{BPComp}(X_i, X_j) \mid \text{base pair } (i,j) \in S \cup S_{\text{PK}_1} \cup S_{\text{PK}_2}\} \tag{2}$$

with $\text{BPComp}(X_i, X_j)$ returns True if $(x_i, x_j) \in \mathcal{BP}$.

**Sequence conservation.** To preserve similarity to natural sequences, positions $i$ exhibiting >90% sequence conservation were restricted to the corresponding set of conserved nucleotides, denoted by $E_i$ (vertex label). This was enforced by adding the constraints

$$\mathcal{C}_{\text{cons}} = \{\text{ValueIn}(X_i, E_i) \mid \text{conserved position } i\} \tag{3}$$

with $\text{ValueIn}(X_i, E_i)$ returning True if $x_i \in E_i$.

**Unique sequence constraint.** Two assignments may result in the same final design as the empty letters are removed. For example, both `A-A` and `-AA` yield the identical sequence `AA` after deletion of empty characters.

To prevent unintended biases during sequence generation, an additional set of constraints $\mathcal{C}_{\text{unique}}$ was imposed to ensure that each designed sequence is uniquely represented by one assignment (with the associated structure). This was enforced by requiring all $\epsilon$ symbols within a region to form a contiguous block located at one end of that region. In other words, once a variable within a region is assigned $\epsilon$, all subsequent variables in the same region are constrained to take the same value. For a helix, the constraint is only required for the 5' strand, since the 3' strand is determined by the base pair complementarity constraints. Thus,

$$\mathcal{C}_{\text{unique}} = \bigcup_{\substack{\text{unpaired or helix 5' strand} \\ \{k,\ldots,k+\ell\}}} \{\text{GapsRight}(X_i, X_{i+1}) \mid \text{position } i \in [k, k+\ell]\} \tag{4}$$

where $\text{GapsRight}(X_i, X_{i+1})$ returns False if $x_i = \epsilon$ and $x_{i+1} \neq \epsilon$. A natural consequence of imposing $\mathcal{C}_{\text{unique}}$ is that the non-empty domain can be assigned immediately to the variables at the beginning of each region while ensuring the minimum size during the base model construction, and can then be omitted from the constraint. The variables on which the constraint GapsRight imposes are marked with a red backbone in Fig. S3.

**Region length correlation constraint.** Analyses of natural xrRNAs also revealed correlations between the lengths of individual structural elements. Although we only considered the most pronounced relationships, i.e., between P3 and L3, into our `Infrared` model, the full set of results can be seen in Fig. S2. Implementing this constraint requires explicit bookkeeping of region lengths for a given assignment, defined as the number of variables in the region that are assigned a non-empty value. Let $X_{i_1}, \ldots, X_{i_r}$ be the variables in the region of interest $R$. We introduce a new set of counter variables $\mathcal{B} = \{B_{i_0}, B_{i_1}, \ldots, B_{i_r}\}$ with the associated domain set $\mathcal{D}_{\mathcal{B}} = \{0\} \times [0, r]^r$. The value of variable $B_{i_0}$ is fixed to $0$, while the rest of variables $B_{i_k}$ take value $b_{i_k}$ from $[0, r]$ representing the amount of non empty values assigned to the variables from $X_{i_1}$ to $X_{i_k}$. The constraint CountValues is then introduced to ensure the correct value is assigned for each $k \in [1, r]$ with

$$\text{CountValues}(B_{i_k}, B_{i_{k-1}}, X_{i_k}) = \text{True if } b_{i_k} = b_{i_{k-1}} + \mathbb{1}_{x_{i_k} \notin \{\epsilon\}}.$$

The last counter variable $B_{i_r}$ represents then the length of the region $R$.

Two types of length correlation are considered in our xrRNA design model:

1. The length of a region $R$ is bounded in a range $[\ell_b, u_b]$ (Tab. S2). The constraint LengthBound is imposed on its last counter variable $B_{i_r}$ with

$$\text{LengthBound}(B_{i_r}, \ell_\beta, u_\beta) = \text{True if } \ell_b \leq b_{i_r} \leq u_b;$$

2. The length ratio of two regions $R$ and $R'$ lies in a desired range $[\ell_b, u_b]$ (Tab. S3). The constraint LengthRatio is imposed on the two last counter variables $B_{i_r}$ and $B'_{i_r}$ with

$$\text{LengthRatio}(B_{i_r}, B'_{i_r}, \ell_\beta, u_\beta) = \text{True if } \ell_b \leq \frac{b_{i_r}}{b'_{i_r}} \leq u_b.$$

The implementation deviates slightly from the formulation presented above in order to avoid redundant computations and reduce run time, without altering the set of valid assignments. In particular, the counter is implemented for the number of the empty letters since less counter variables are needed.

**Table S2:** Regions where the total length is controlled using additional constraints.

| Subclass | Regions | Length Range |
|---|---|---|
| MBFV | P2 & L2 | $[7, 15]$ |
| | P3 & L3 | $[20, 23]$ |
| | L3 | $[5, 15]$ |

**Table S3:** Regions where the length ratio is controlled using additional constraints.

| Subclass | $R$ | $R'$ | Ratio Range |
|---|---|---|---|
| MBFV | P3 | L3 | $[0.8, 1.2]$ |

We have introduced the needed variables, domains, and constraints to define the sequence space of the given symbolic representation. An example of such a sequence satisfying all constraints is given in Fig. S3 B. The additional variables used to count the individual regions lengths are placed next to the corresponding base positions. Instead of uniformly sampling the sequence space, we consider in this study a Boltzmann weighted sampling provided by `Infrared` with the sequence weight defined by the following two features.

**Total length feature.** To sample the sequence with lengths similar to natural sequences ($54 \pm 7$ nt), we introduced a length feature counting the number of non-empty assignments

$$F_{len} = \{\text{NotEmpty}(i) \mid \text{position } i \in \{1, ..., n\}\}. \tag{5}$$

The feature is composed of network functions NotEmpty$(i)$, which return $1$ if $X_i \neq \epsilon$ and $0$ otherwise. The feature evaluation of an assignment is then the designed sequence length.

**Minimum free energy feature.** The second feature is used to influence the folding energy of the target structure towards the values found in nature (-18 kcal/mol). It has been shown in [5] that a simple base pair energy model is sufficient to target the Turner energies in sequence sampling. We therefore introduce two features, associated with different weights, controlling the energy of the base nested structure and pseudoknot PK2:

$$F_{\text{energy}} = \{\text{BPEnergy}(i, j) \mid (i, j) \in S\} \tag{6}$$

$$F_{\text{PKenergy}} = \{\text{BPEnergy}(i, j) \mid (i, j) \in S_{PK2}\} \tag{7}$$

Here, the function BPEnergy$(i, j)$ decomposes the structure energy into the local contribution of each base pair $(X_i, X_j)$ as described in [5].

The appropriate weights for each feature were determined before the post-sampling optimization. Sequences were sampled while changing the weights until the target value is reached. Those weights were then fixed during the optimization.

**Optimization with Monte Carlo.** To improve sequence specificity toward the target xrRNA fold, we performed post-sampling optimization using a Monte Carlo approach as suggested in [15]. The optimization procedure encompasses two stages. First, the ensemble defect was minimized using a sampled sequence as the starting point. The resulting optimum was subsequently used to initialize a second stage in which the target frequency was maximized.

This two-stage strategy was found to be effective because the ensemble defect is particularly sensitive to changes far from the optimum and tends to favor sequences where misfoldings still remain structurally competent. Maximizing the target frequency in the second stage then refines the sequence such that the target structure becomes the minimum free-energy (MFE) fold and competing conformations are energetically disfavored, while retaining the benefit of a low ensemble defect.

Throughout the process, the Monte-Carlo temperature was fixed to $T = 0.01$, and the objective functions were calculated based on the pseudoknot-free target structure using `ViennaRNA` [7]. A total of 250 000 optimization steps, equally split between the two objective functions, were performed. At each step, one connected component of the dependency network was selected with a probability relative to its size, and the variables of this connected component were reassigned with new values according to the model described above. At step $t$, a new assignment with an objective value of $O_t$ was accepted with a probability

$$p(O_t, O_{t-1}) = \begin{cases} 1 & \text{if } O_t \geq O_{t-1} \\ e^{\frac{O_t - O_{t-1}}{T}} & \text{if } O_t < O_{t-1} \end{cases} \tag{8}$$

where $T$ is the Monte-Carlo temperature. For objectives defined on positive real values, such as the ensemble defect, minimization was implemented by maximizing the corresponding negative objective value.

### D *In silico* function validation

The *in silico* validation aimed to quantify the force required to disrupt the ring-like fold of the candidate designs. These force profiles were compared against a natural xrRNA with established XRN1 resistance to benchmark candidate performance.

The initial tertiary structures were created using `SimRNA` [1] and `QRNAS` [10] as described in the Methods section. Molecular dynamics (MD) systems were prepared with the implicit-solvent builder in `Charmm-GUI` [6, 11] using the GBn2 implicit solvent model optimized for nucleic acids [9]. Simulations were carried out at implicitly simulated 0.15 mol/L salt concentration and 300 K employing the AMBER OL3 force field [16]. Each system was simulated for 500 ns to obtain an equilibrated ensemble. Representative conformations were extracted and rigidly transformed such that the 5′ end was positioned at the origin of the coordinate system.

Steered molecular dynamics (SMD) simulations were performed in `OpenMM` [4] without a cutoff for nonbonded interactions and using hydrogen bond constraints. To simulate the exoribonuclease tension on the 5' end of the xrRNA, an external pulling force was applied in the $z$ direction to the most 5' phosphor atom, using a linear gradient of 1000 pN/µs. While this force gradient exceeds physiological conditions, it enables efficient probing of the stability of the ring-like structure without the computational cost of resolving the full unfolding pathway.

The XRN1 active site was approximated using a membrane in the (x/y)-plane containing a pore with a diameter of 12 Å centered at the origin. This boundary was implemented via a custom external potential based on the repulsive component of a Lennard-Jones interaction. To prevent initial lateral drift away from the pore, additional restraints were applied to confine the 5′ end near the pore axis. Four independent replicas were simulated using a Langevin integrator with the default timestep of 2 fs and friction coefficient of 1 $\text{ps}^{-1}$. The pulling force applied to the xrRNA was updated every 1000 steps according to the force gradient, starting from 0 pN at the first time step.

Because the present analysis focused on the force required to overcome the initial mechanical barrier associated with the ring-like topology, simulations were terminated once the 5′ end had translocated more than 120 Å through the pore (typically after 500–1000 ns). Consequently, the trajectories are not intended to resolve the complete unfolding mechanism, but rather provide an efficient compromise between computational cost and a quantitative stability readout.

### E xrRNA Design Sequences

**Table S4:** Sequences of xrRNA substrates for *in vitro* exonuleolytic degradation tests.

xrRNA_wt

5’-GCUAAAAAGACACCUACCGGUGUCAAAGUCAGGCCAGAAAUGCCACCGGAUAAAGGUAGACGGUGCUGCCUGCAACCUUUCUGCGGAAGGAAUAACCGCAGU-3’

△PK1

5’-GCUAAAAAGACACCUACCGGUGUCAACCUCAGGCCAGAAAUGCCACCGGAUAAAGGUAGACGGUGCCGCCUGCAACCUUU-3’

PK2_4bp

5’-GCUAAAAAGACACCUACCGGUGUCAAAGUCAGGCCAGAAAUGCCACCGGAUAUGCUUAGACGGUGCUGCCUGCAACGUAU-3’

PK2_2bp

5’-GCUAAAAAGACACCUACCGGUGUCAAAGUCAGGCCAGAAAUGCCACCGGAUAUGUGUAGACGGUGCUGCCUGCAAAAUCU-3’

PK2_1bp

5’-GCUAAAAAGACACCUACCGGUGUCAAAGUCAGGCCAGAAAUGCCACCGGAUAGAAGUAGACGGUGCUGCCUGCAAGUAGU-3’

△PK2

5’-GCUAAAAAGACACCUACCGGUGUCAAAGUCAGGCCAGAAAUGCCACCGGAUCGAAAGAGACGGUGCUGCCUGCAAAAAAA-3’

syn-xrRNA1

5’-GCUAAAGAGCUUGCGUAAAGCAAGCAAAGUCAGGCCGGGUCCCCCGCCACGUGGAGCCCUUAUCCGCGUGCUGCCUGUAGGGAA-3’

syn-xrRNA2

5’-GCUAAAGAGCUUGCGUAAAGCAAGCAAAGUCAGGCCGGGUCCCCCGCCACGUGGAGCCCCCAUCCGCGUGCUGCCUGCCGGGGGA-3’

syn-xrRNA3

5’-GCUAAAGCCGCAUAGAGCGGCAAAGUCGGCCGUACGAAUAGUACCCAGGACAGCCUCCCUCUGUCCUGCUGGCCGUGGGAGAG-3’

### F Evaluation of XRN1 stop sites

To identify XRN1 stop sites within the xrRNA constructs, incomplete degradation products were sequenced and mapped to the corresponding reference xrRNA sequence. Stop positions were inferred by plotting the frequency of read-start coordinates along the reference. A distinct peak marks the predominant XRN1 stop site. Additional peaks at the first position indicate that untreated xrRNAs have been sequenced. The relative abundance of full-length xrRNA and degradation products cannot be used to quantify protection efficiency in this assay.

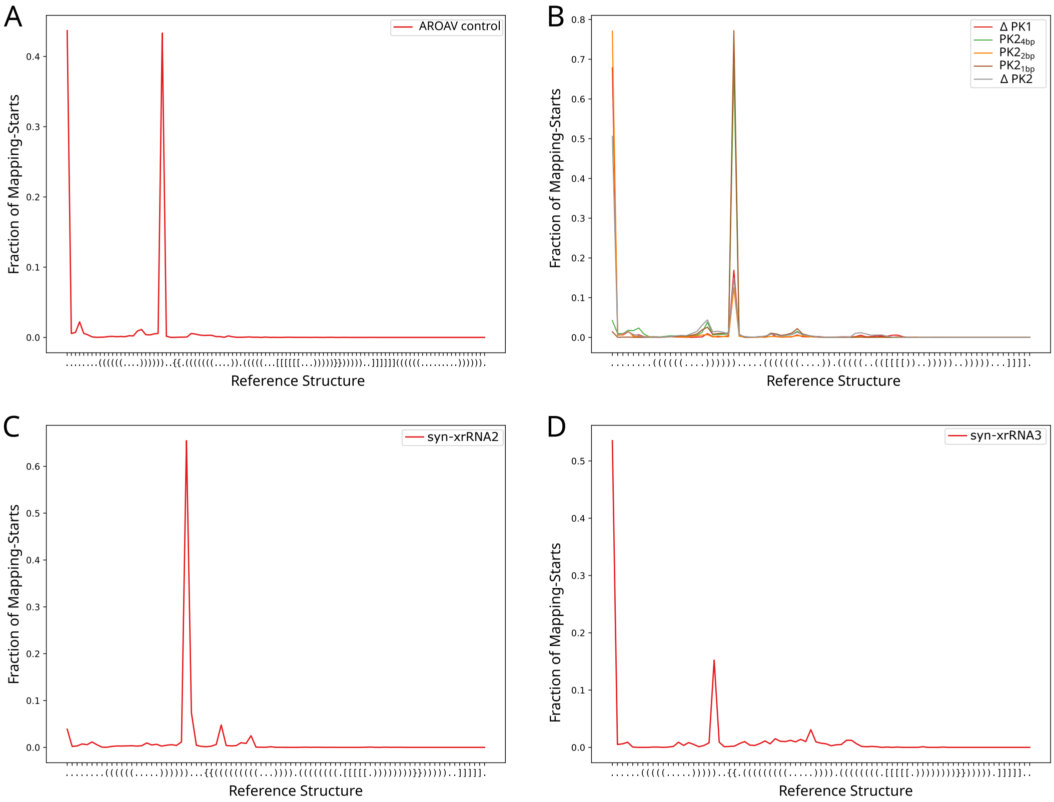


**Figure S4: XRN1 Stop point analysis.** Frequency of read-start positions obtained from sequencing of incomplete degradation products after XRN1 incubation for AROAV wild-type xrRNA (A), mutants with altered PK1 and PK2 stability as described in Fig. 2 (B), syn-xrRNA2 (C), and syn-xrRNA3 (D).

#### G XRN1 degradation of AROAV-xrRNA mutants

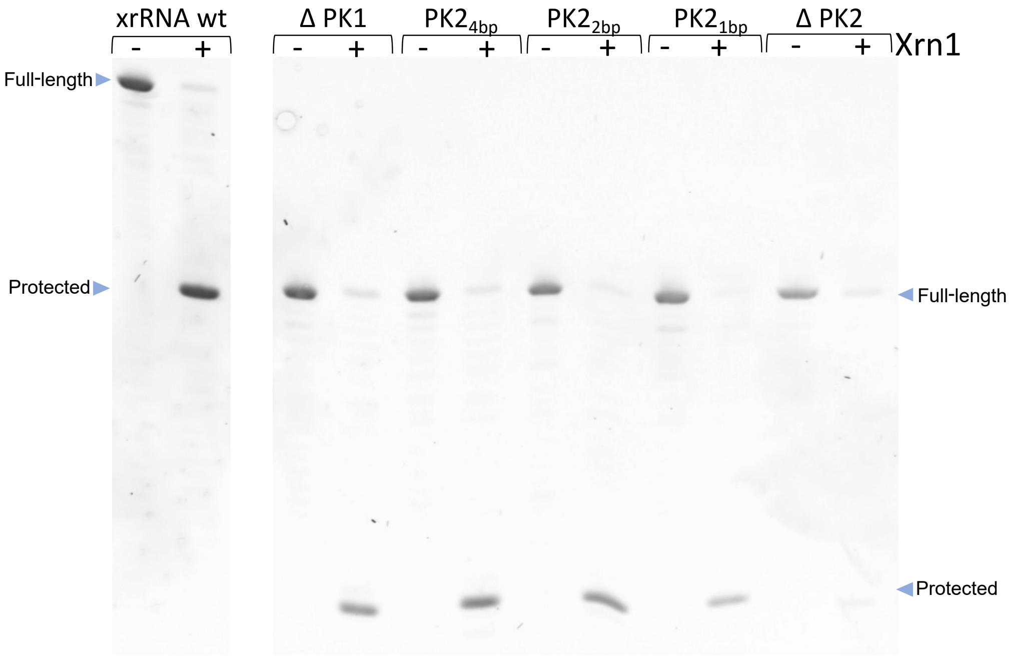


**Figure S5: XRN1 degradation of xrRNA mutants with varying PK stability.** Degradation assay of the reference AROAV wild-type xrRNA (left) and mutants with systematically altered pseudoknot stability (right). Mutant constructs were derived from the wild-type sequence by deleting PK1 and PK2, or by progressively weakening PK2 through stepwise reduction of the number of base pairs.

#### H Molecular dynamics simulations of synthetic xrRNA designs

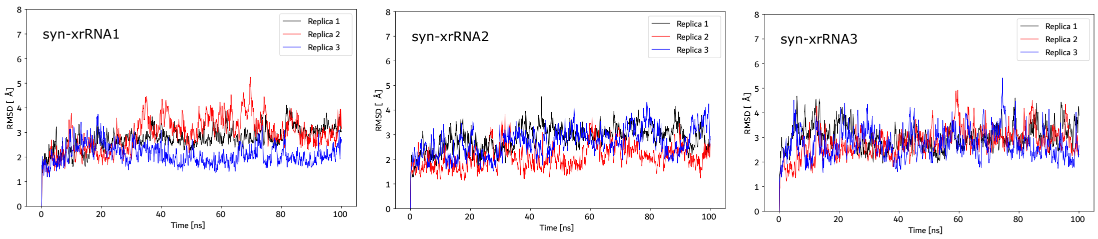


**Figure S6: Stability of xrRNA designs in MD simulations.** Root mean square deviation (RMSD) over 100 ns for three independent explicit-solvent MD replicas of the xrRNA designs following energy minimization and equilibration.

### I Structure probing of designed xrRNAs

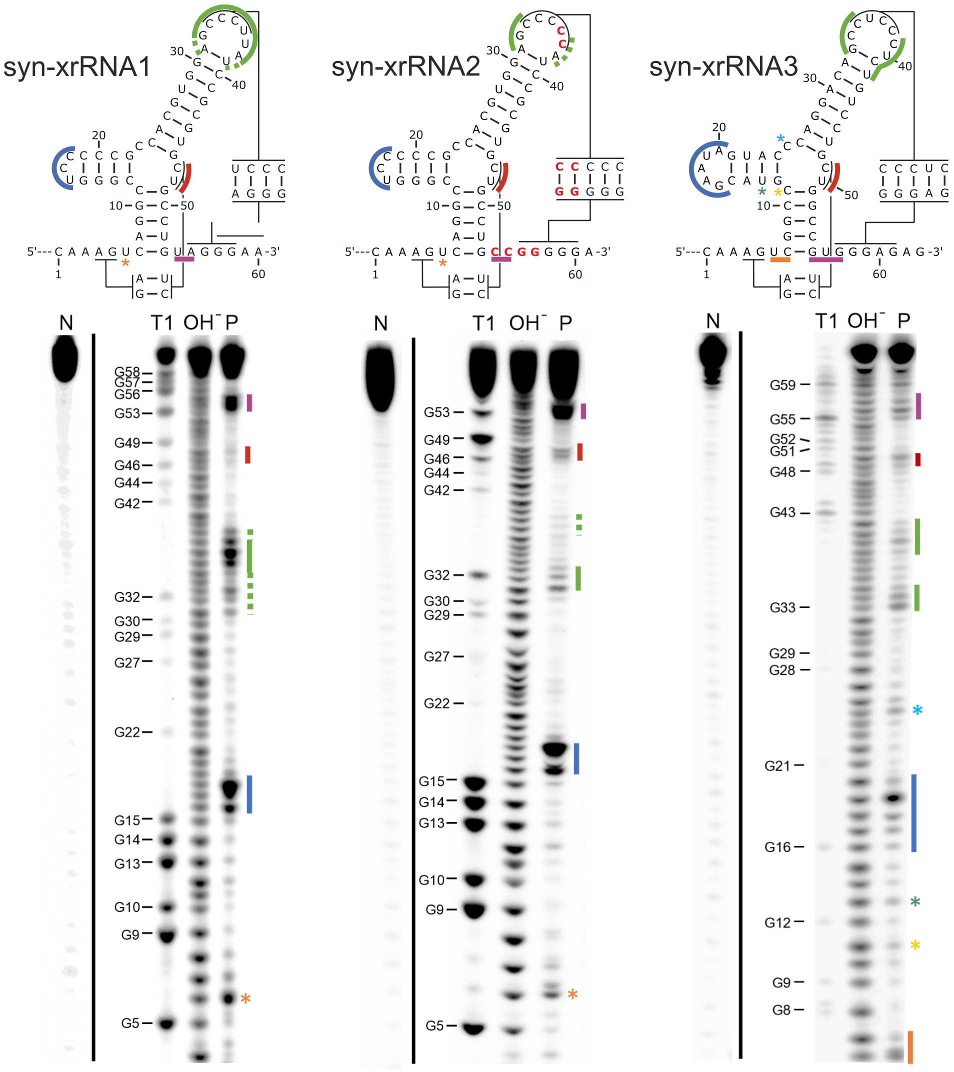


**Figure S7: In-line probing of syn-xrRNA 1 to 3.** In the secondary structure schematics, cleavage-accessible regions are indicated by colored bars (or asterisks for single nucleotide positions), with colors corresponding to the labeled band patterns in the autoradiographs. While syn-xrRNA1 exhibits partial fraying in the loop region involved in PK2 formation, this region is clearly base-paired in syn-xrRNA2 and syn-xrRNA3. By contrast, the downstream segment of PK1 (sequence CU located in the 3-way junction) shows detectable cleavage in all three constructs, indicating that PK1 is less stable than PK2 and, consequently, contributes less to the mechanical robustness of the protective ring-like structure. This is consistent with the observation that deletion of PK2 does not fully abolish exonuclease resistance (Fig. 3). N, negative control; T1, G-specific cleavage by T1 nuclease (numbering according to the depcisted secondary structures); OH-, alkaline hydrolysis of RNA samples; P, in-line probing.

#### J Novelty of xrRNA designs

To assess whether our designs are substantially diverged from the biological sequences used to parameterize the `Infrared` model, we created a covariance model from the natural MBFV xrRNAs (MBFV-CM). The designs were then evaluated against this model and, in addition, against the Rfam covariance model for xrRNAs (RF03547-CM), see Tab. S5. The trusted cutoff for RF03547-CM is 40.2 as noted in Rfam. None of the designs exceeded this threshold, indicating that they are not classified as xrRNAs with high confidence by the general Rfam model. This observation is consistent with limited representation of MBFV xrRNAs in the training set underlying RF03547-CM.

As expected, MBFV-CM readily detected the first two designs, which still retain substantial sequence similarity to natural xrRNAs. In contrast, syn-xrRNA3, generated with most sequence constraints removed, and driven primarily by structural constraints, was not recognized by the MBFV-CM, indicating that the sequence is further removed from the biological sequence space captured by the covariance model.

An alternative to the rational `Infrared`-based workflow is to sample new sequences from a covariance model using `cmemit`. The differences between these methods are visualized in Fig. S8. As can be seen from the base pairing probability matrix of the designed sequences, `cmemit`-generated sequences do not reliably reproduce the pseudoknot interactions and exhibit substantially higher propensity for alternative base pairings, consistent with an increased risk for misfolding.

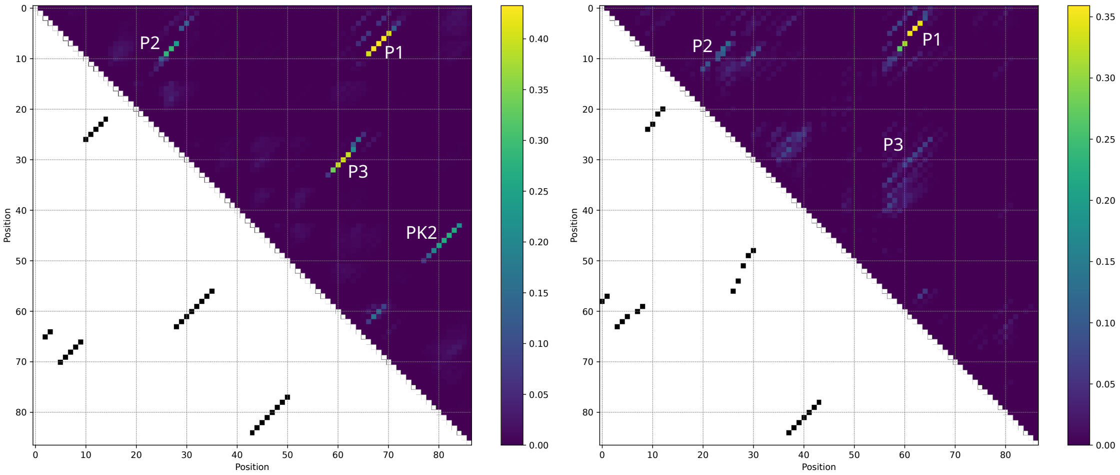


**Figure S8: Comparison between `Infrared` and `cmemit` designs.** Base-pairing probability matrices for sequences designed with the `Infrared` model (left) and the MBFV covariance model (right). Probabilities were obtained by averaging base pairing probabilities across all designs within each set. The upper triangles show the averaged pairing probabilities with annotated structural elements (P1, P2, P3, PK2), whereas the lower triangles show the corresponding target structure.

**Table S5:** Similarity of xrRNA designs relative to natural sequences. A database comprising all four designs was searched against different covariance models: the Rfam model for general xrRNAs (RF03547-CM), and a covariance model constructed from the sequences used to derive the `Infrared` constraints (MBFV-CM). No Rfam covariance model specific to mosquito-borne flavivirus xrRNAs was available. None of the designs exceeded the noise cutoff for the Rfam models. For MBFV-CM, the two highest-scoring hits are highlighted.

| Design | RF03547-CM | | MBFV-CM | |
|---|---|---|---|---|
| | score | E-value | score | E-value |
| syn-xrRNA1 | 5.0 | 0.27 | **25.9** | **5.0e-09** |
| syn-xrRNA2 | 5.9 | 0.18 | **23.0** | **4.9-09** |
| syn-xrRNA3 | 4.2 | 0.38 | 5.1 | 0.06 |

[3] Karen Clark, Ilene Karsch-Mizrachi, David J Lipman, James Ostell, and Eric W Sayers. Genbank. *Nucleic Acids Res*, 44(D1):D67–D72, 2016.
